# Democratizing computational pathology: optimized Whole Slide Image representations for The Cancer Genome Atlas

**DOI:** 10.1101/2023.12.04.569894

**Authors:** Tristan Lazard, Marvin Lerousseau, Sophie Gardrat, Anne Vincent-Salomon, Marc-Henri Stern, Manuel Rodrigues, Etienne Decencière, Thomas Walter

## Abstract

Automatic analysis of hematoxylin and eosin (H&E) stained Whole Slide Images (WSI) bears great promise for computer assisted diagnosis and biomarker discovery. However, scarcity of annotated datasets leads to underperforming models. Furthermore, the size and complexity of the image data limit their integration into bioinformatic workflows and thus their adoption by the bioinformatics community. Here, we present Giga-SSL, a self-supervised method for learning WSI representations without any annotation. We show that applying a simple linear classifier on the Giga-SSL representations improves classification performance over the fully supervised alternative on five benchmarked tasks and across different datasets. Moreover, we observe a substantial performance increase for small datasets (average gain of 7 AUC point) and a doubling of the number of mutations predictable from WSIs in a pan-cancer setting (from 45 to 93). We make the WSI representations available, compressing the TCGA-FFPE images from 12TB to 23MB and enabling fast analysis on a laptop CPU. We hope this resource will facilitate multimodal data integration in order to analyze WSI in their genomic and transcriptomic context.

---

Cancer diagnosis heavily relies the examination of H&E stained tissue slides, which offer crucial insights into the disease and potential treatment options and which are routinely acquired in pathology labs. Digitizing these slides into Whole Slide Images (WSI) enables automated analysis, aiming at assisting clinicians in executing tedious tasks, such as counting mitoses [1], identification of metastases[2] and grading[3]. Furthermore, the availability of large data repositories, such as The Cancer Genome Atlas (TCGA) provides us with the challenging opportunity to identify morphological biomarkers related to survival[4] or treatment response[5], and to unravel the complex genotype-phenotype relationships by building predictive models for molecular features, such as single gene mutations[6], mutational signatures[7] and molecular subtypes [6].

However, WSI are not yet extensively used outside the pathology community for two primary reasons. First, the size and complexity of WSI require special skills and equipment for their analysis. A single WSI may contain billions of pixels and thousands of cells, complicating storage, processing, and analysis. Second, while pathology labs are generating ever increasing WSI datasets, annotated WSI datasets are often scarce, in particular for rare diseases, specific molecular subtypes or in the context of clinical trials. Training current deep learning models on such datasets often leads to underperforming models with poor generalization capability.

Self-supervised learning offers a promising approach for addressing these chalenges. This training paradigm leverages unlabeled datasets to pretrain neural networks which then demonstrate improved performance when fine-tuned on smaller, annotated datasets. Numerous studies in the computational pathology field have already adopted self-supervised learning, but only at the tile level ([3, 7–9]). They used such pretrained networks to encode the small images that compose the WSI-the tiles-, effectively reducing them from billions of pixels to a few thousand feature vectors. These feature vectors then serve as the basis for training multiple instance learning (MIL) algorithms ([9–12]). Nonetheless, the sheer volume of tiles per WSIs still makes MIL models both computationally intensive to train and susceptible to overfitting. While is a fair effort toward training without supervision wide histopathological images -up to 4096 pixel squared-, they do not succeed in training WSI representations.

Here, we introduce Giga-SSL, a novel self-supervised method that utilizes large, unlabeled WSI datasets to learn compact and highly discriminative WSI-level features. It can encode a WSI into a single vector of 512 values, and we show that a simple logistic regression operating on these representations achieves equal or better performance than fully-supervised MIL architectures across several tasks and datasets. Furthermore, we open-source these representations for the entire TCGAformalin-fixed, paraffin-embedded (FFPE), reducing its size from 12 TB to 23 MB without loss of predictive power.

## Results

### Giga-SSL provides a framework for self-supervised learning at the slide-level

For this, Giga-SSL models a WSI as an array of tile representations, with the objective to apply contrastive learning (CL) on this array. CL is a SSL framework whose main task is to bring together the representations of two randomly transformed versions of the same object, while pushing away -*contrasting* the representations of different objects.

In order to optimize this objective at the scale of WSIs, we devised a specialized approach that involves both tile-scale and slide-scale transformations. This design is executed through a two-step architecture, as illustrated in Figure 1A. The architecture consists of two distinct neural networks; the first network, which is pretrained on histopathological images, is responsible for encoding the transformed tiles, and only the second network, comprising sparse convolutional layers, undergoes optimization during the giga-SSL training process.

**Fig. 1.**
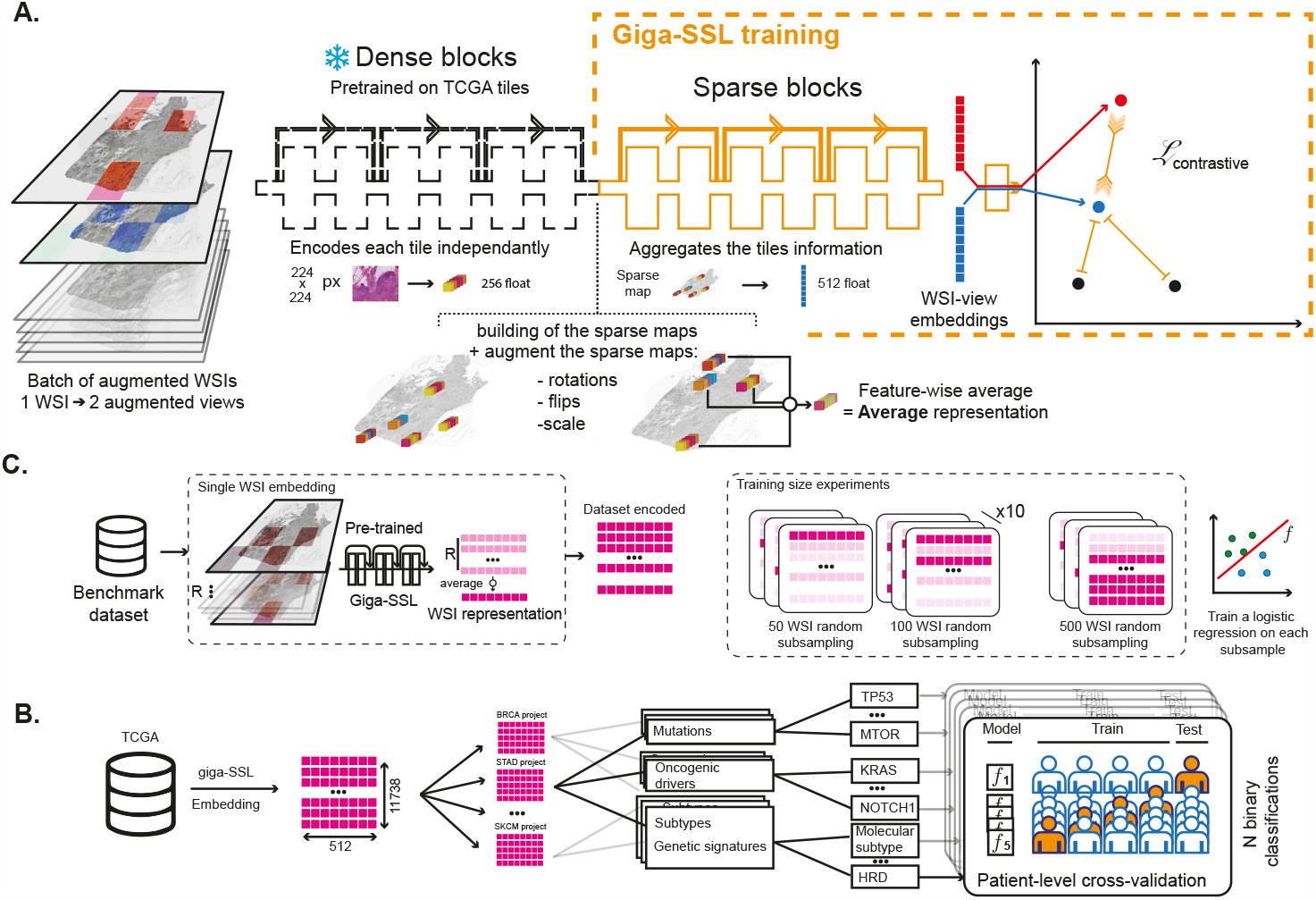
**A**. Overview of the Giga-SSL architecture and training procedure: Initially, two distinct views of the same Whole Slide Image (WSI) are created using tile and slide level augmentations. Each augmented tile is encoded using a pre-trained convolutional tile-encoder. The objective of Giga-SSL training is to optimize the second sparse convolutional block to minimize a contrastive loss at the slide level. **B**. Benchmark tasks experiments workflow, detailing the computation of a WSI representation. **C**. Workflow of the pancancer classification experiments.

The output of this second block, the WSI representations, are used in all the downstream analysis tasks by training L2-regularized logistic regressions. For the sake of brevity, we refer to these models as *Giga-SSL classification models*. Their corresponding performance metrics are designated as *Giga-SSL performances*.

### Giga-SSL outperforms fully-supervised methods on several classification tasks and across cancer types

We first compared Giga-SSL models to state-of-the-art WSI classification algorithms across five TCGA benchmark tasks. These include breast cancer subtyping (lobular/ductal), lung cancer subtyping (lung adenocarcinoma (LUAD)/ lung squamous cell carcinoma (LUSC)), kidney subtyping (clear/papillary/chromophobe cells), and two breast cancer-related tasks: homologous recombination deficiency / proficiency (HRD/HRP) and molecular profiling (Triple Negative Breast Carcinoma (TNBC)/luminal). We gradually reduced the training dataset size through stratified subsampling down to 50 WSIs and assessed the performance of models trained on these subsets (see Fig. 1B). We compared Giga-SSL to the CLAM-SB algorithm [10] operating on the same tile representations as the one used by the Giga-SSL model. Fig. 2 **A**. shows the performance difference between the Giga-SSL and CLAM models as a function of the training set size. Each point above the red line indicates superior performance of Giga-SSL. Across all tasks, using 100% of the training data, Giga-SSL consistently achieves state-of-the-art performance. The advantage of Giga-SSL over CLAM grows as the training set size shrinks. With only 50 WSIs, Giga-SSL provides an average gain of 7 AUC points over CLAM.

**Fig. 2.**
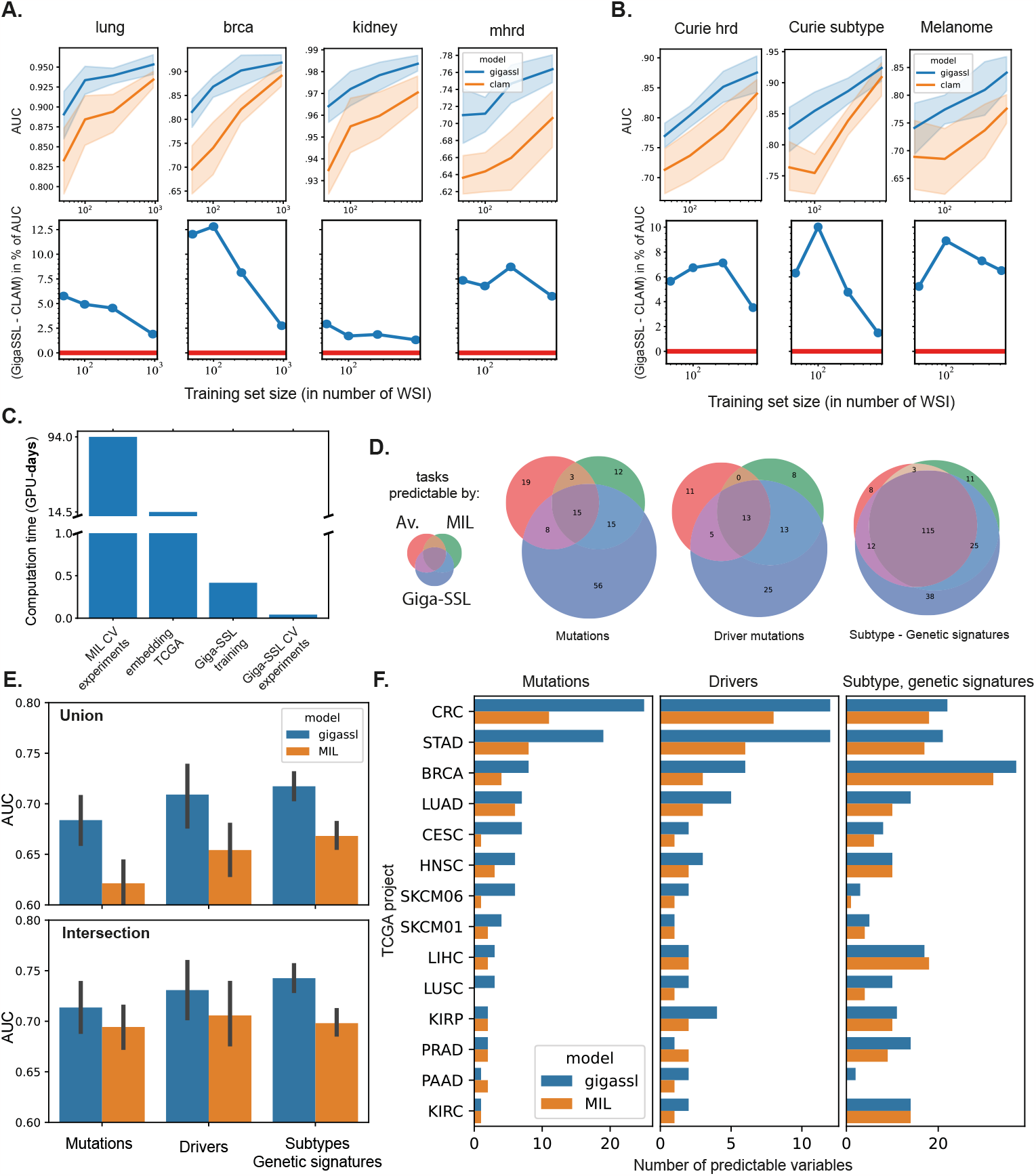
Evaluation Results. **A**. Displays the results for the four benchmark tasks. The first row of graphs presents the ROC-AUC scores of Giga-SSL and CLAM as a function of the training set size. The second row demonstrates the AUC improvement of Giga-SSL over CLAM as a function of the training set size. **B**. Presents the results on the external validation datasets for both Giga-SSL and CLAM. **C**. Indicates the time requirements for key computation steps, measured in GPU-days. CV: Cross-validation. **D**. The number of predictable tasks for each model and each task type (mutations, driver mutations, subtypes, and genetic signatures) in a pancancer setting. **E**. Displays the average cross-validated AUC for the three different types of classification tasksmutation, oncogenic drivers, subtypes and genetic signatures-for Giga-SSL and MIL models. In the upper panel, the results are averaged across all tasks that are predictable by the Giga-SSL **or** MIL model (union). The lower one shows averages across the tasks predictable by both models. Error bars are indicate standard deviations of these distributions of AUC. **F**. Provides a detailed breakdown of item D., focusing on the granularity of the TCGA projects.

**Fig. 3.**
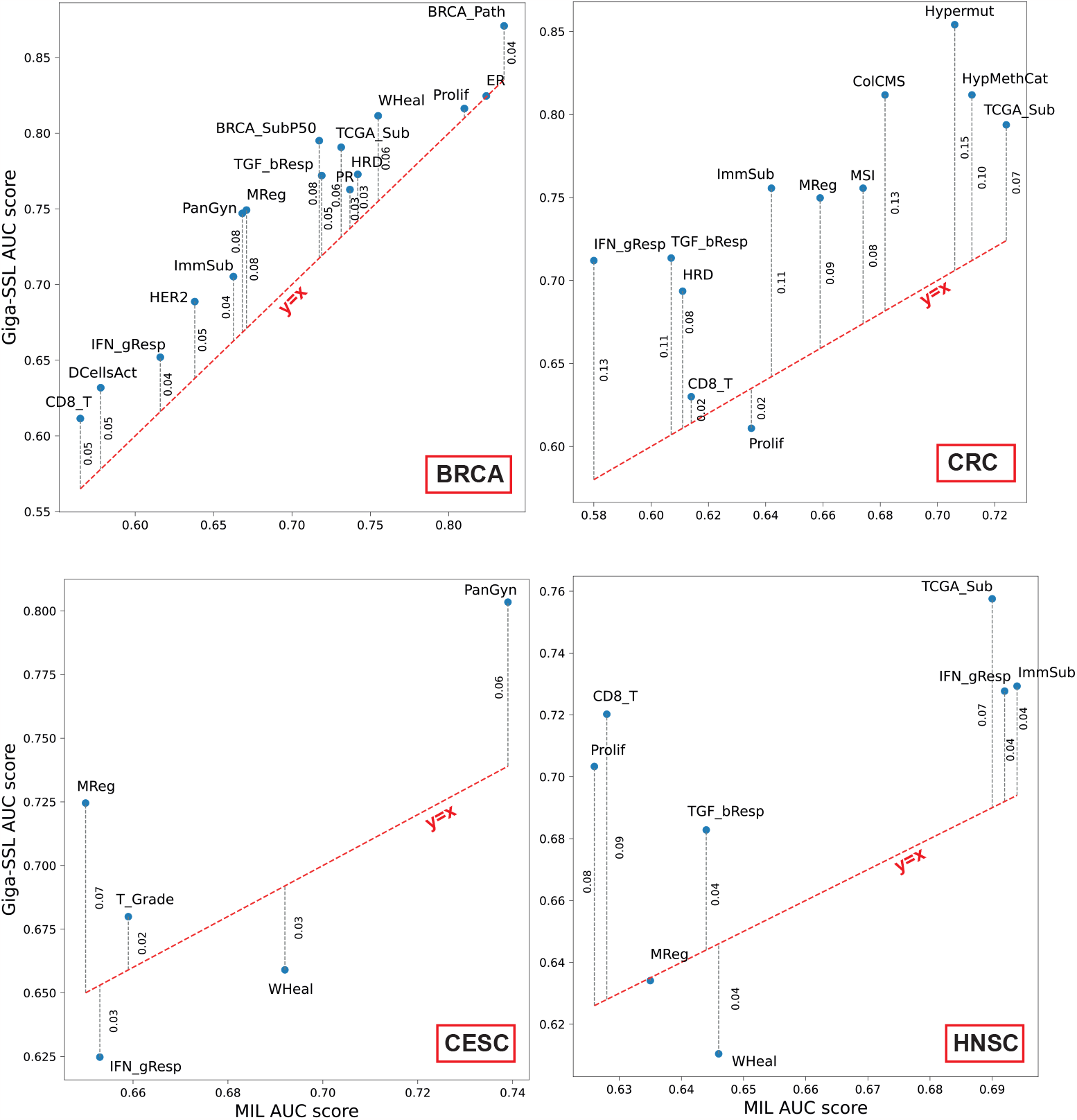
Performances of the non-mutation tasks for BRCA, CRC, CESC and HNSC TCGA projects. Plots shows the performances of the logistic regression trained on top of the Giga-SSL features against the performances of the MIL model used in [6]. Red line indicates same performances between the two models. PR - PRStatus (Progesterone Receptor); AR - AR protein (Androgen Receptor); N HistGrade - Neoplasm Histologic Grade; T Grade - Tumor Grade; DCellsAct - Activated Dendritic Cells; SCNA - Somatic Copy Number Alterations; TCGA Sub - TCGA Subtypes; HER2 - HER2 Final Status; ImmSub - Immune Subtypes; HypMethCat - Hypermethylation Category; WHeal - Wound Healing; CD8 T - T Cells CD8; PanGyn - Pan-Gynecologic Clusters; GHistClass - Gastric Histological Classification; GrowthPat - Major Growth Pattern; Prolif - Proliferation; ClinGleason - Clinical Gleason Sum; PanKidPath - Pan-Kidney Pathology; IFN gResp - IFN-gamma Response; MReg - Macrophage Regu- lation; HomRecDef - Homologous Recombination Defects; HCCSub - Hepatocellular Carcinoma Subtypes; ERG - ERG Status; ColCMS - Colorectal Cancer CMS; MSI - Microsatellite Instability Status; ER - Estrogen Receptor Status; BRCA Path - BRCA Pathology; Hypermut - Hypermutated; TGF bResp - TGF-beta Response; BRCA SubP50 - BRCA Subtype (PAM50)

**Fig. 4.**
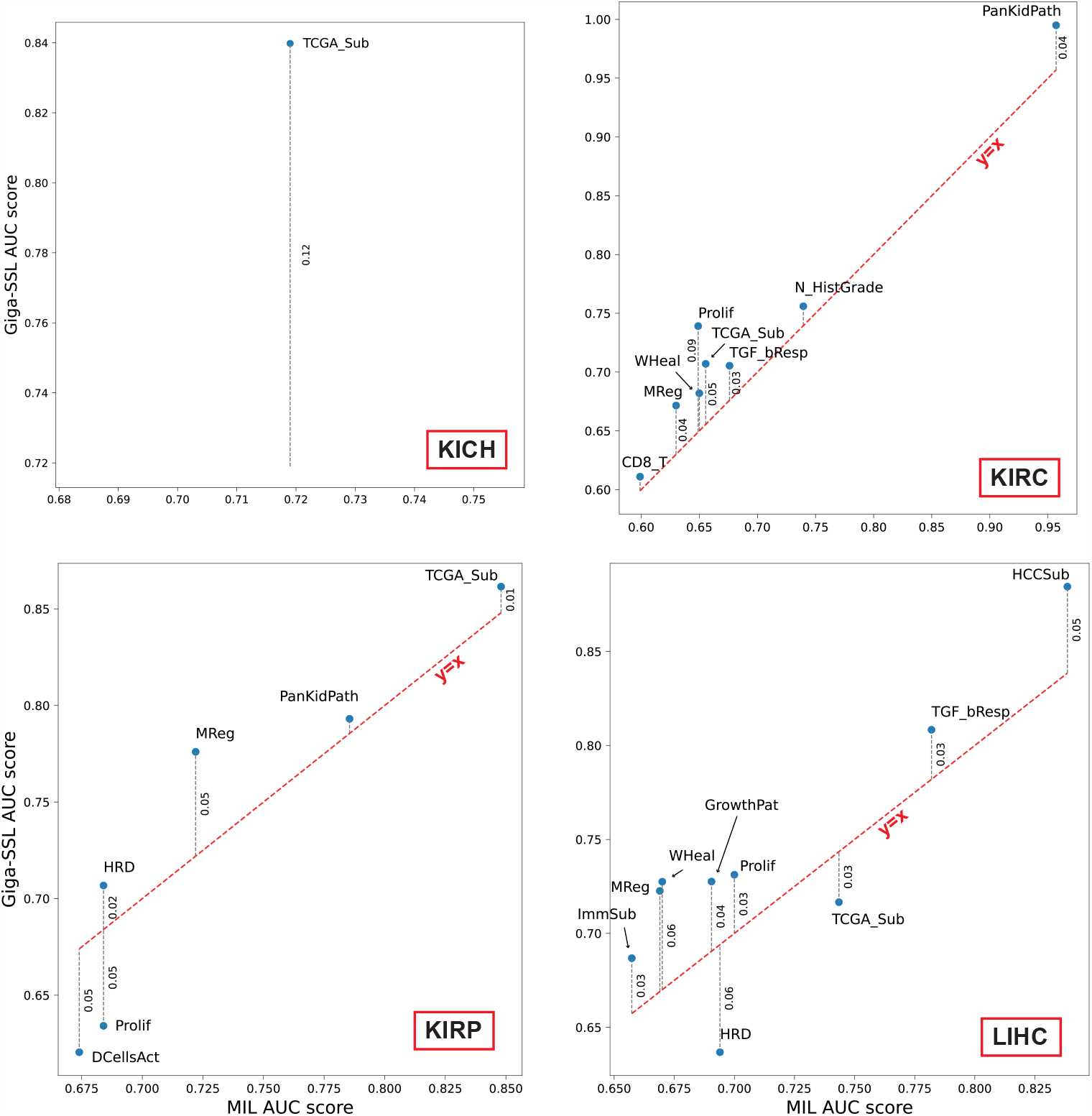
Performances of the non-mutation tasks for KICH, KIRC, KIRP and LIHC TCGA projects. Plots shows the performances of the logistic regression trained on top of the Giga-SSL features against the performances of the MIL model used in [6]. Red line indicates same performances between the two models. PR - PRStatus (Progesterone Receptor); AR - AR protein (Androgen Recep- tor); N HistGrade - Neoplasm Histologic Grade; T Grade - Tumor Grade; DCellsAct - Activated Dendritic Cells; SCNA - Somatic Copy Number Alterations; TCGA Sub - TCGA Subtypes; HER2 - HER2 Final Status; ImmSub - Immune Subtypes; HypMethCat - Hypermethylation Category; WHeal - Wound Healing; CD8 T - T Cells CD8; PanGyn - Pan-Gynecologic Clus- ters; GHistClass - Gastric Histological Classification; GrowthPat - Major Growth Pattern; Prolif - Proliferation; ClinGleason - Clinical Gleason Sum; PanKidPath - Pan-Kidney Pathology; IFN gResp - IFN-gamma Response; MReg - Macrophage Regulation; HomRecDef - Homologous Recombination Defects; HCCSub - Hepatocellular Carcinoma Subtypes; ERG - ERG Status; ColCMS - Colorectal Cancer CMS; MSI - Microsatellite Instability Status; ER - Estrogen Receptor Status; BRCA Path - BRCA Pathology; Hypermut - Hypermutated; TGF bResp - TGF-beta Response; BRCA SubP50 - BRCA Subtype (PAM50)

**Fig. 5.**
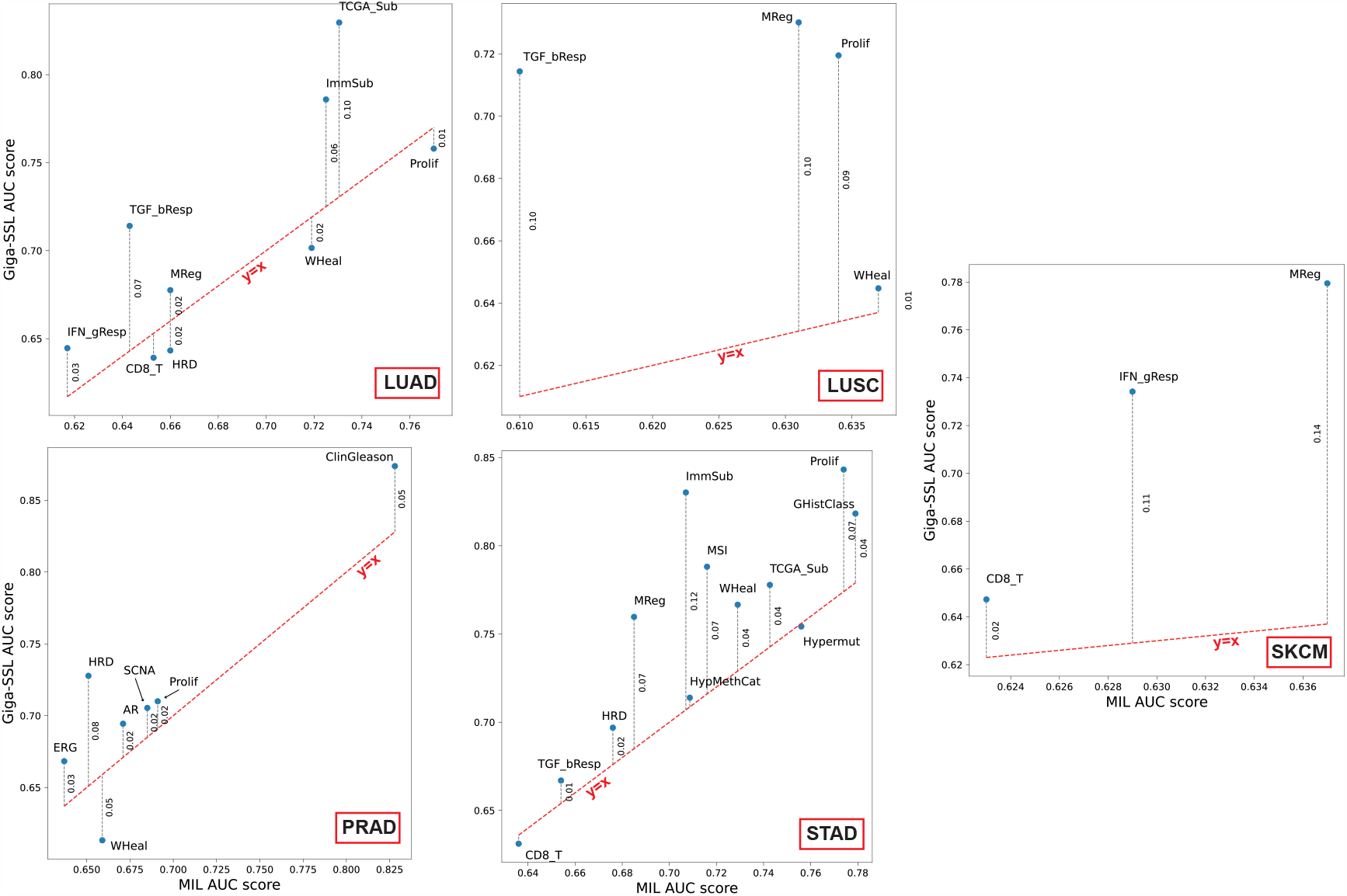
Performances of the non-mutation tasks for LUAD, LUSC, PRAD, STAD and SKCM TCGA projects. Plots shows the performances of the logistic regression trained on top of the Giga-SSL features against the performances of the MIL model used in [6]. Red line indicates same performances between the two models. PR - PRStatus (Progesterone Receptor); AR - AR protein (Androgen Recep- tor); N HistGrade - Neoplasm Histologic Grade; T Grade - Tumor Grade; DCellsAct - Activated Dendritic Cells; SCNA - Somatic Copy Number Alterations; TCGA Sub - TCGA Subtypes; HER2 - HER2 Final Status; ImmSub - Immune Subtypes; HypMethCat - Hypermethylation Category; WHeal - Wound Healing; CD8 T - T Cells CD8; PanGyn - Pan-Gynecologic Clusters; GHistClass - Gastric Histological Classification; GrowthPat - Major Growth Pattern; Prolif - Proliferation; ClinGleason - Clinical Gleason Sum; PanKidPath - Pan-Kidney Pathology; IFN gResp - IFN-gamma Response; MReg - Macrophage Regu- lation; HomRecDef - Homologous Recombination Defects; HCCSub - Hepatocellular Carcinoma Subtypes; ERG - ERG Status; ColCMS - Colorectal Cancer CMS; MSI - Microsatellite Instability Status; ER - Estrogen Receptor Status; BRCA Path - BRCA Pathology; Hypermut - Hypermutated; TGF bResp - TGF-beta Response; BRCA SubP50 - BRCA Subtype (PAM50)

### Giga-SSL transfers well to other datasets

To confirm these findings, we performed external validation experiments to explore the generalization capabilities of our representations. For this we applied Giga-SSL to two in-house WSI datasets for two different cancer types:

- Breast cancer (BC): 788 in-house H&E stained WSIs from BC patients with known Homologous Recombination status. We conducted two classification tasks: HRD prediction and subtype prediction (TNBC/luminal) [7].
- Uveal Melanoma (UM): 516 in-house H&E stained WSIs from UM patients. The objective here was to predict chromosome 3 status (disomy 3 or monosomy 3, which represent two major UM subtypes with contrasted prognosis) [14].

Fig. 2.B shows the improved performance of logistic regression with Giga-SSL representations compared to CLAM. The average AUC increase is 3.8% using 100% of the data and 5.7% using 50 WSIs: Giga-SSL weights maintain their performance and label efficiency when applied to unseen datasets.

### Giga-SSL can predict more mutations and genetic signatures than MIL

We next turned to the prediction of mutations and genetic signatures across cancer types. Kather et al. showed in a seminal study that many mutations and genetic signatures are predictable from H&E stained tissue slides[6]. Our objective was to assess whether logistic regression could effectively predict mutations and signatures using Giga-SSL representations, as an alternative to employing a full MIL. The data workflow for these experiments is delineated in Fig. 1D. In total, the prediction tasks are (1) 830 point mutation predictions across 14 tumor types (2) 376 known oncogenic driver mutations and (3) 182 subtyping and genetic signature tasks. Of note, genetic variant prediction tasks are usually imbalanced, with the minority class comprising 8% of the dataset on average. This contrasts with subtyping and genetic signature prediction tasks, where the minority class represents on average 33% of the dataset. The Venn diagram in Figure Fig. 2.D illustrates the number of mutations that can be predicted by the Giga-SSL model (represented by the blue areas), the Average model -see methods and Fig. 1A.- (shown as the red area), and the MIL model [6] (depicted by the green area). We observe that Giga-SSL is able to predict most of the mutations that are predictable with the previously published method (30/45 for the point mutations, 26/34 for the oncogenic driver mutations). Furthermore, the Giga-SSL representations allow the prediction of an additional 64 point mutations and 30 driver mutations, which roughly doubles the number of predictable mutations from WSIs across these 14 cancer types. Details about the predictable mutations are available in Supplementary Tables 3-6. A similar result is obtained for genetic signatures: out of 374 binary classification tasks (see methods) the majority of signatures predictable by the MIL model[6] can also be predicted by Giga-SSL (140 out of 154) and the average model (127 out of 154). Compared to the MIL model, the Giga-SSL model can predict an additional 50 signatures. Additionally, as illustrated in Fig. 2E, Giga-SSL provides enhanced classification accuracy across all tasks. Details about the performances of Giga-SSL on these tasks are available in Supplementary Figures 3-5

### Giga-SSL is modular

Contrary to monolithic frameworks and algorithms, Giga-SSL relies on the correct adjustment of different training elements: the tile-encoder pre-training, tile-level and slide-level augmentations, and the aggregation module. Each module can be updated independently, allowing the entire framework to continuously benefit from advancements in their respective domains. We illustrate this by comparing various pre-trained tile-encoders as shown in Supplementary Table 1, notably the recent ctranspath network, and demonstrate that the WSI Giga-SSL representation benefits from an improved tile-encoder; a research effort that already receives much attention [15–17].

### Giga-SSL is computationally efficient

To highlight Giga-SSL’s computational efficiency, Fig. 2.C shows the GPU-days required for our experiments. Training a Giga- SSL model for 1000 epochs on the complete TCGA dataset takes only 10 GPU-hours on a single GPU. Once we have obtained the representations, downstream prediction tasks become highly efficient. This is because the classification algorithm fits only logistic regression, and the cross-validated experiments take approximately 1 hour on a laptop CPU — a sharp contrast to the 92 GPU-days reported in [6]. When applied to external datasets, Giga-SSL showcases good scalability. On average, a WSI can be encoded by Giga-SSL in roughly 5 seconds using a single GPU. This implies that encoding a typical dataset of 1000 WSIs can be completed in less than 1.5 hours.

## Discussion

In this article, we present a framework providing generic representations for H&E stained WSI, targeting robust solutions for small datasets and simplified training, thus lowering barriers for morpho-molecular cancer analyses. Addressing these challenges, we introduce Giga-SSL, the first SSL framework able to derive concise yet highly discriminative WSI-level embeddings.

Using logistic regression, we highlight the advantages of these features, achieving state-of-the-art classification performances on small datasets, exhibiting robust generalization to external datasets and doubling the number of predictable gene mutations in the TCGA across cancer types.

A major advantage of Giga-SSL is its computational efficiency, at training and inference time. This efficiency allows for the potential training of Giga-SSL models on datasets significantly larger than the TCGA at minimal cost and with much lower environmental impact. Moreover, even scientists without detailed knowledge of image analysis and deep learning can easily utilize this modality, facilitating tests on new outcome variables or experiments with different datasets stratifications. We are releasing the entire encoded TCGA-FFPE dataset to the public, reducing its size from 12 TB to 23 MB. Such readily available embeddings can be integrated into TCGA genomics and transcriptomic analyses without any need for prior image-processing know-how or specialized equipment. We are optimistic that this initiative will spark interest within the bioinformatics sector, encouraging comprehensive integration of pathology and molecular data, and fostering joint exploration of cancer’s molecular and morphological landscape.

## Methods

### Tile embeddings

We obtain tile embeddings using contrastive learning. Specifically, we employ MoCo [18], training a ResNet18 on 6 million tiles extracted from a random set of 3000 FFPE slides from the TCGA over 200 epochs using MoCoV2’s standard transformation. The tile embeddings are obtained through spatial pooling of the activations of the third block of this network. The tile representations are kept static, meaning they are not further optimized during the Giga-SSL training phase.

### Giga-SSL architecture

The architecture of Giga-SSL combines the ResNet18 framework with SparseConvMIL [19]. The first four residual convolution blocks independently encode each tile. The resulting tile encodings are aggregated within a SparseMap and processed by 4 sparse convolution blocks [19]. Together, these components achieve functionality akin to a full ResNet18, as detailed in [20], and the final layer of this architectural blocks are the WSI embeddings used in the downstream analysis. A final multi-layer perceptron called *projector* further encodes the embeddings of the WSI, following [18]. The CL loss is computed using the output of the projector network.

#### Slide-level transformations

aim to generate different views that are to be pulled closer or pushed apart, depending on whether they originate from the same WSI, while maintaining part of the biological information of the WSI. To generate these views, we use the following transformations:

- Subsampling: Tiles are randomly sampled among all the tissue-tiles of a WSI. The harshness of the transformation is given by the number *T* of tiles subsampled per view. Notably, when *T* = 1, the Giga-SSL training framework becomes equivalent to HiDisc-Slide [21].
- Tile transformations: The tiles are randomly augmented with classical image augmentations (hue, rotation, Gaussian blur, flips, crops) before being encoded by the tile encoder. Training Giga-SSL requires this transformation to be *shared* among views; that is, when building a transformed view of a WSI, the same augmentation must be applied to all sampled tiles.
- Sparse-map transformations: The sparse-maps undergo scale, rotation and flips transformation, augmenting the geometries of the WSIs.

### Giga-SSL Training

Training is performed in 2 steps. First, the tile encoder is pre-trained on histopathology data using MoCo [18] and frozen, like in [20]. This allows to compare Giga-SSL to competing methods based on the same tile encodings. In a second step, the sparse units (in orange 1) are trained on the full TCGA with the WSI-level contrastive pretext task under the slide-level transformations mentioned in the text. Training is performed for 100 GPU-hours on a single V100 GPU. We use the Adam optimizer with a starting learning rate of 0.003. We use a cosine annealing learning rate scheduler with 10 warming epochs and a final learning rate to 3*e*^*−*6^. As described in the methodological publication [20], we approximate the tile-level augmentations by randomly sampling 25 augmentations per WSI. We then uniformly sample 64 tiles per WSI per augmentation, augment and embed them using the tile-encoder described previously.

### WSI representation computation

To regularize the embeddings, the Giga-SSL network is applied to *R* = 50 different views of each slide. The Giga-SSL representations are the average of these 50 runs. The Average representations are the feature-wise average of the representations of all the tiles of a WSI. We call *Average models* the logistic regressions operating on these representations. Both the Average and Giga-SSL representation are then unit-normalized using scikit-learn Normalizer.

### Statistical procedure

To address downstream classification challenges, we deploy an L2-regularized logistic regression (parameters: C=10, class weight=‘balanced’).

We report the performance of these logistic regressions over 10 random dataset splits for benchmarking and generalization experiments, and 3 splits for mutation prediction experiments (to account for the very imbalanced nature of the mutation prediction task). The same splits are used in compared methods.

For mutation prediction, the primary criterion is task predictability. As outlined in [6], we accumulate the model’s posterior probabilities predicted throughout the crossvalidation folds. We then conduct a two-sided t-test between the predictions of the samples belonging to one class and the predictions from the remainder of the dataset. We adjusted the resulting p-values for multiple testing, accounting for a total of 1388 classification tasks, using the Benjamini-Hochberg correction. We set the significance threshold, *p*thres, to 0.01 and compare the predictability of tasks between Giga-SSL and the MIL model implemented in the original publication. Each task of the pancancer experiments employs a patient-level three-fold cross validation strategy.

## Data Availability

The TCGA-FFPE WSIs are publicly available through https://portal.gdc.cancer.gov/. The in-house Curie datasets (breast cancer slides and UV) consist of confidential medical data not open to the public.

The lightweight embeddings of all the TCGA-FFPE slides computed with giga-SSL models trained using MoCo and CtransPath embeddings, as well as the used models, are made publicly available at https://data.mendeley.com/datasets/d573xfd9fg/3 (DOI: 10.17632/d573xfd9fg.3).

Labels for various datasets are extracted from different sources:

- Lung (LUAD/LUSC): GDC Portal
- Breast Cancer (Ductal/Lobular): GDC Portal
- Kidney (clear/papillary/chromophobe cells): GDC Portal
- mhrd Breast Cancer (HRP/HRD): GerkeLab Repository, as in [22]. HRD score are binarized using mean as threshold.
- All pancancer experiments: Supplementaries of [6]. The continuous variables are binarized using mean as threshold.

## Code Availability

We open-source the following softwares:

- A python package designed to train Giga-SSL models : https://github.com/trislaz/gigassl.
- A Python package for encoding WSIs using Giga-SSL models is available at https://github.com/trislaz/democratizing_WSI. The package was designed to be userfriendly, enabling a wide range of users to utilize GigaSSL and our pre-trained models.

## Acknowledgements

This work was performed using HPC resources from GENCI-IDRIS (Grant 2023-AD011011863R3). The results in this article are based upon data generated by the TCGA Research Network: https://www.cancer.gov/tcga. TL was supported by a Q-Life PhD fellowship (Q-life ANR-17-CONV-0005). Furthermore, this work was supported by the French government under management of Agence Nationale de la Recherche as part of the “Investissements d’avenir” program, reference ANR-19-P3IA-0001 (PRAIRIE 3IA Institute) and by the ITMO Cancer (20CM107-00).

## A Supplementaries

**Table 1.** Effect of the tile-encoder on Giga-SSL representation performance. Results show a 5-fold patient-level stratified cross-validated average. Models are logistic regressions (C=10, class weight=‘balanced’) built on Giga-SSL representations using various tile-encoders. Advances in tile-encoder design enhance the efficacy of Giga-SSL representations. The tasks referenced are the 5 benchmark tasks from the main paper. The same cross-validation folds are used for all the experiment of the same task.

| Tile-encoder | task | AUC | F1 Score | Balanced Accuracy |
| --- | --- | --- | --- | --- |
| MoCo [18] | brca | $0.919 \pm 0.023$ | $0.792 \pm 0.024$ | $0.84 \pm 0.031$ |
| | kidney | $0.982 \pm 0.014$ | $0.891 \pm 0.033$ | $0.91 \pm 0.028$ |
| | lung | $0.959 \pm 0.014$ | $0.897 \pm 0.012$ | $0.898 \pm 0.012$ |
| | mhrd | $0.783 \pm 0.042$ | $0.734 \pm 0.035$ | $0.734 \pm 0.035$ |
| | thrd | $0.843 \pm 0.032$ | $0.778 \pm 0.036$ | $0.779 \pm 0.036$ |
| CtransPath [16] | brca | $0.95 \pm 0.024$ | $0.849 \pm 0.019$ | $0.875 \pm 0.026$ |
| | kidney | $0.991 \pm 0.005$ | $0.929 \pm 0.023$ | $0.943 \pm 0.016$ |
| | lung | $0.968 \pm 0.013$ | $0.914 \pm 0.023$ | $0.914 \pm 0.022$ |
| | mhrd | $0.814 \pm 0.045$ | $0.749 \pm 0.032$ | $0.749 \pm 0.032$ |
| | thrd | $0.879 \pm 0.026$ | $0.795 \pm 0.036$ | $0.796 \pm 0.036$ |

**Table 2.** Detail of the composition of the datasets used for the benchmark tasks.

|  | N Patients | N Slides | Labels repartition |
| --- | --- | --- | --- |
| TCGA - brca | 1041 | 977 | 831 (ductal) / 210 (lobular) |
| TCGA - lung | 1033 | 936 | 528 (TCGA-LUAD) / 505 (TCGA-LUSC) |
| TCGA - mhrd | 912 | 853 | 465 (0) / 447 (1) |
| TCGA - kidney | 924 | 882 | 510 (ClearCell) / 294 (Papillary) / 120 (Chromophobe) |
| TCGA - thrd | 634 | 586 | 318 (HRD) / 316 (HRP) |
| Curie - HRD | 787 | 787 | 485 (HRD) / 302 (HRP) |
| Curie - Subtype | 787 | 787 | 514 (luminal) / 273 (TNBC) |
| Curie Melanoma | 515 | 515 | 336 (Monosomy) / 179 (Disomy) |

**Table 3.** All point mutations predictable by Giga-SSL and the MIL of [6], sorted by TCGA project.

| Project | Mutations predictables by GigaSSL and MIL |  |  |  |  |
| --- | --- | --- | --- | --- | --- |
| **BRCA** | MAP3K1 | PIK3CA | TP53 |  |  |
| **CESC** | STK11 |  |  |  |  |
| **CRC** | APC | BRAF | KMT2B | KMT2D | KRAS |
|  | MGA | PIK3CA | PTCH1 | RNF43 | TP53 |
| **HNSC** | CASP8 | NSD1 | TP53 |  |  |
| **KIRC** | PBRM1 |  |  |  |  |
| **KIRP** | SETD2 |  |  |  |  |
| **LIHC** | CTNNB1 |  |  |  |  |
| **LUAD** | PDGFRB | TP53 |  |  |  |
| **PRAD** | TP53 |  |  |  |  |
| **STAD** | FBXW7 | KMT2B | KMT2C | KMT2D | MTOR |
|  | PIK3CA | TP53 |  |  |  |

**Table 4.** All point mutations newly predictable with the Giga-SSL features, sorted by TCGA project.

| Project | Mutations predictables by GigaSSL only |  |  |  |  |  |
| --- | --- | --- | --- | --- | --- | --- |
| **BRCA** | BRCA1 | ERBB2 | FBXW7 | GATA3 | PBRM1 |  |
| **CESC** | APC | ERBB2 | KMT2D | KRAS | TCERG1 | TP53 |
| **CRC** | ACVR2A | ATM | BRCA2 | CDC27 | FAT1 | FBXW7 |
|  | JAK2 | MIER3 | MSH6 | PPP6C | PTEN | RB1 |
|  | RHOA | TCERG1 | TTN |  |  |  |
| **HNSC** | APC | HRAS | JAK2 |  |  |  |
| **KIRP** | PBRM1 |  |  |  |  |  |
| **LIHC** | TP53 | TSC2 |  |  |  |  |
| **LUAD** | EGFR | KRAS | MED12 | NFE2L2 | TTN |  |
| **LUSC** | KEAP1 | RAC1 | TP53 |  |  |  |
| **PAAD** | TP53 |  |  |  |  |  |
| **PRAD** | TTN |  |  |  |  |  |
| **STAD** | ACVR2A | ARHGAP6 | ARID1A | B2M | CDK12 | KMT2A |
|  | MAP3K1 | MET | MGA | RBM10 | RHOA | TCERG1 |
| **SKCM01** | MGA | PIK3CA | RNF43 | TTN |  |  |
| **SKCM06** | CDC27 | CDKN2A | FBXW7 | GNAS | KDM6A | PPP6C |

**Table 5.** Oncogenic driver mutations predictable by Giga-SSL and the MIL of [6], sorted by TCGA project.

| **Project** | Driver mutations predictable by GigaSSL and MIL |  |  |  |  |
| --- | --- | --- | --- | --- | --- |
| **BRCA** | MAP3K1 | PIK3CA | TP53 |  |  |
| **CRC** | APC | BRAF | KMT2B | KMT2D | KRAS |
|  | PIK3CA | RNF43 | TP53 |  |  |
| **HNSC** | NSD1 | TP53 |  |  |  |
| **KIRC** | PBRM1 |  |  |  |  |
| **KIRP** | KRAS | SETD2 |  |  |  |
| **LIHC** | CTNNB1 |  |  |  |  |
| **LUAD** | EGFR | TP53 |  |  |  |
| **PAAD** | KRAS |  |  |  |  |
| **PRAD** | TP53 |  |  |  |  |
| **STAD** | BRCA2 | FBXW7 | KMT2B | KMT2D | PIK3CA |

**Table 6.** Oncogenic driver mutations newly predictable with the Giga-SSL features, sorted by TCGA project.

| **Project** | Driver mutations predictable only by GigaSSL |  |  |  |
| --- | --- | --- | --- | --- |
| **BRCA** | BRCA1 | GATA3 | SMAD4 |  |
| **CESC** | KRAS | STK11 |  |  |
| **CRC** | BRCA2 | MGA | NRAS | PTEN |
| **HNSC** | CASP8 |  |  |  |
| **KIRC** | TP53 |  |  |  |
| **KIRP** | MET | PBRM1 |  |  |
| **LIHC** | TP53 |  |  |  |
| **LUAD** | KRAS | MGA | U2AF1 |  |
| **LUSC** | NFE2L2 | PIK3CA |  |  |
| **PAAD** | TP53 |  |  |  |
| **SKCM01** | PIK3R1 |  |  |  |
| **SKCM06** | CDKN2A | CTNNB1 |  |  |
| **STAD** | AMER1 | ARID1A | KMT2C | PTCH1 |
|  | RHOA | RNF43 | TP53 |  |

## Notes

https://data.mendeley.com/datasets/d573xfd9fg/3

### Competing Interest Statement

The authors have declared no competing interest.

